# Genetic and Immunological Evidence for Microbial Transfer Between the International Space Station and an Astronaut

**DOI:** 10.1101/2020.11.10.376954

**Authors:** David C. Danko, Nitin Singh, Daniel J. Butler, Christopher Mozsary, Peng Jiang, Ali Keshavarzian, Mark Maienschein-Cline, George Chlipala, Ebrahim Afshinnekoo, Daniela Bezdan, Francine Garrett-Bakelman, Stefan J. Green, Fred W. Turek, Martha Hotz Vitaterna, Kasthuri Venkateswaran, Christopher E. Mason

## Abstract

Microbial transfer from the environment can influence a person’s health, but relevant studies often have confounding variables and short durations. Here, we used the unique environment of the International Space Station (ISS) to track movement of microbes between an astronaut’s commensal microbiomes and their environment. We identified several microbial taxa, including *Serratia proteamaculans* and *Rickettsia australis* which appear to have been transferred from the ISS to the commensal microbiomes of the astronaut. Strains were matched at the SNP and haplotype-level, and notably some strains persisted even after the astronaut’s return to Earth. Some transferred taxa correspond to secondary strains in the ISS environment, suggesting that transfer may be mediated by evolutionary selection. Finally, we show evidence that the T-Cell repertoire of the astronaut changes to become more specific to environmental taxa, suggesting that continual microbial and immune monitoring can help guide spaceflight mission planning, health monitoring, and habitat design.

## 1 Introduction

Human commensal microbiomes have a known hereditary component (Goodrich et al., 2016), but the non-hereditary, acquired portion of the human microbiome is still being defined in terms of its covariates and components. It is known that the microbiome can change as a function of age, diet, developmental stage, environmental exposures, antibiotic use, and lifestyle, yet strain-level mapping and longitudinal tracking of such dynamics are limited. In particular, the movement of non-pathogenic microbes and how they can colonize an adult commensal microbiome in a defined, quantified, and sealed environment, is almost completely unknown (Schwendner et al., 2017). An ideal study for microbial transfer would utilize a longitudinal sampling of subjects in a hermetically-sealed environment that was already profiled with strain-level resolution.

As an environment, the International Space Station (ISS) presents several key advantages for the study of microbial transfer. It is a well studied environment, with microbial tracking studies ongoing since 2014, and its occupants’ microbiomes are routinely sequenced. Moreover, the ISS is a uniquely sealed environment with essentially no chance of infiltration by exterior microbes, save for the regular supply missions. Finally, microgravity on the ISS may lead to an improved diffusion of microorganisms relative to studies done in terrestrial environments.

Evidence for the transfer environmental microbes into adult commensal microbiomes could have important health implications, as it would provide a mechanism for how regional environmental microbiomes impact a person’s microbiome. Cities in particular are known to host diverse environmental microbiomes (Danko et al., 2019) and transfer between commensal and environmental microbiomes may add to explanations for health differences between otherwise similar regions (Nicolaou et al., 2005). The selective transfer of certain microbial strains may also carry evolutionary implications for the microbes being transferred. If a microbial species can be shown to follow distinct selective patterns inside and outside of human commensal microbiomes, or even on the ISS, it is possible that these patterns could guide strain or even species differentiation.

Here, we present genetic and immunological evidence for the transfer of environmental strains to an astronaut’s gut and oral microbiome while on the International Space Station (ISS) during an almost year-long mission(Garrett-Bakelman et al., 2019a). The strain-level data was compared to the T-cell receptor (TCR) diversity and sequence changes during the mission, and these increased matches midflight corresponded to the candidate microbial strains observed. Of note, several of these strains were still observed for months after the mission, providing evidence of a persistent influence on the astronaut’s microbiome, which may help to inform future studies on human microbial interaction.

## 2 Results

We collected 18 fecal and 23 oral microbiome samples from two identical twin human astronauts, one flight subject (TW, 9 stool, 6 saliva, 5 buccal) and one control who did not leave Earth (HR, 9 stool, 7 saliva, 5 buccal), taken from 2014-2018. These were compared to 42 time-matched, environmental samples from the ISS that corresponded to the flight subject’s mission duration. All samples were sequenced with 2×150bp read length to a mean depth of 12-15M reads (12.01, 14.96, and 14.97M mean reads for ISS, fecal, and saliva, respectively), then aligned to the catalog of NCBI RefSeq complete microbial genomes, examined for single nucleotide polymorphisms (SNPs), and then run with strain analysis with the MetaSUB CAP pipeline and Aldex2 (see Methods).

### 2.1 Taxonomic profiles show evidence of continual microbial exchange

#### New taxa in flight subject (TW) match environmental and commensal microbiomes

We first examined the proportion of taxa observed in a given sample that were not observed in a previous sample from the same donor. Any newly observed taxa in sample of a given type (e.g. stool) was annotated relative its presence in samples from other body or environmental sites (e.g saliva). For fecal samples, we segmented the previously unobserved taxa from each sample into four groups: taxa observed in any saliva sample taken before the given fecal samples, taxa observed in ISS samples but not observed in saliva, taxa observed in both ISS and saliva samples, and taxa that were not observed in either the ISS or the saliva. The same process was repeated for saliva samples but swapping fecal and saliva in the hierarchy. As expected,the time series of samples taken from the flight subject (TW) and ground control subject (HR) showed that earlier samples exhibited a greater proportion of novel organisms (Figure 1, S1).

**Figure 1:**
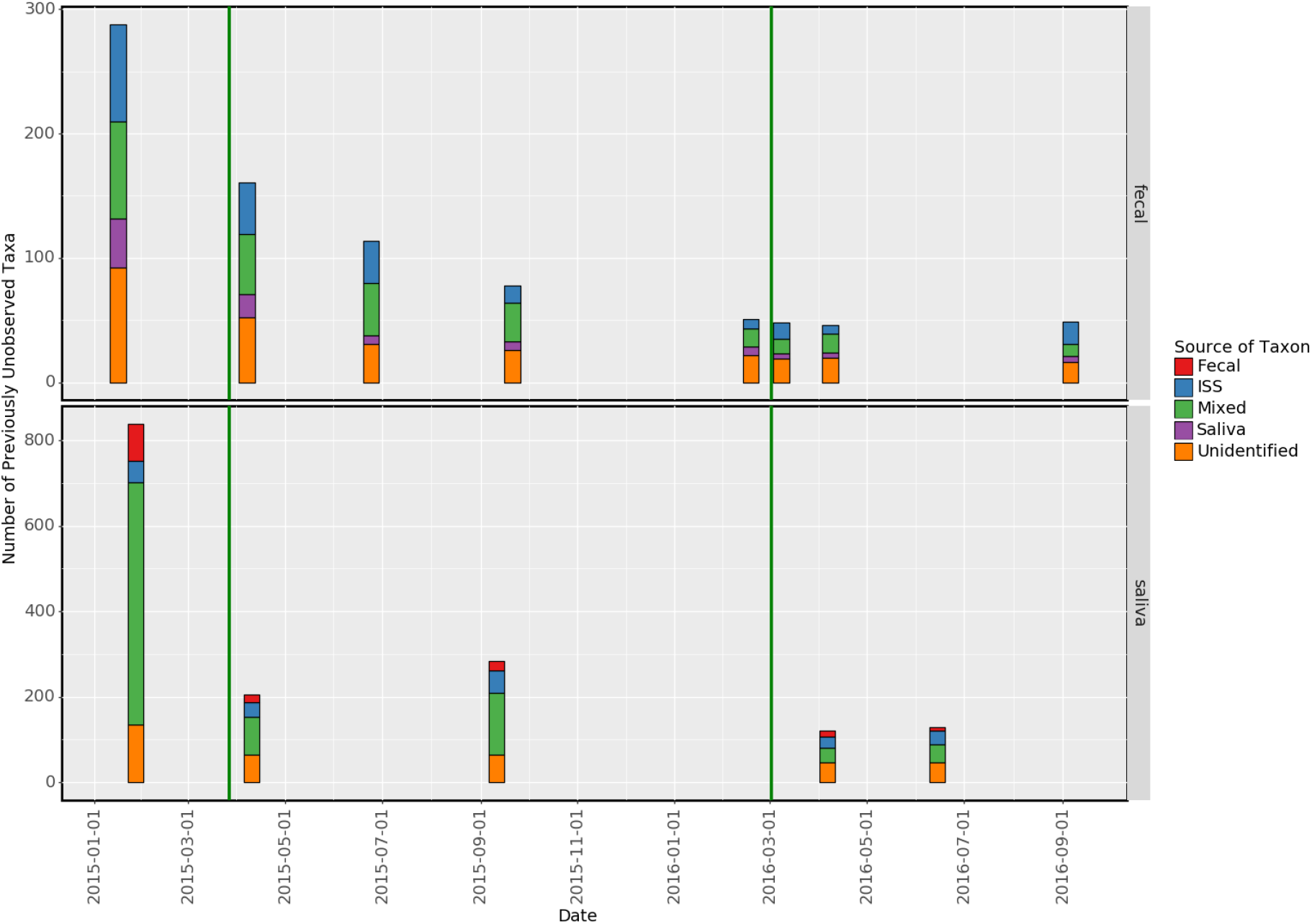
This plot shows the number of taxa at each time point that were not observed at any previous timepoint for fecal and saliva samples from TW. The colors indicate the likely source of the new taxon if it was found previously in the saliva (for fecal samples, vice versa for saliva samples), the ISS, both (Mixed), or neither.

Of note, each sample contains a number of unobserved taxa that matched taxa from saliva/feces or the ISS (even before flight), indicating these are common commensal species on Earth or possibly organisms absorbed in previous missions. Indeed, both astronauts had previously been in the space station across multiple missions though with a 10-fold difference in duration (TW has logged 520 total days on the ISS vs. 54 days for HR). Interestingly, when we examined the fraction of taxa that match ISS taxa in pre-flight samples from TW compared to other samples from HR, a higher average rate (56%) of ISS-matching taxa was observed in pre-flight samples for TW relative to HR (51%), although not significant (p-value = 0.21). The fraction of taxa that matched different environments are listed in Table 1. For both saliva and fecal microbiomes, the large majority of taxa at each time point had already been observed in a previous sample from that site.

**Table 1:** This table gives the average percent overlap between the number of emergent taxa in fecal and saliva microbiomes and microbiomes in other sites.

| Commensal Type | Fecal | Saliva |
| --- | --- | --- |
| Sites Where Taxa Originated |  |  |
| Fecal Only | n/a | 8.7 |
| Saliva Only | 10.2 | n/a |
| ISS Only | 24.5 | 17.6 |
| Both ISS & Saliva/Fecal | 29.9 | 44.6 |
| Taxa not identified in another site | 35.5 | 29.1 |

A small number of taxa were never observed in any pre-flight sample from any body site from either HR or TW, but were observed in mid- and post-flight samples from TW. We thus filtered for taxa that had no reads observed in pre-flight samples and had at least ten reads in at least two mid- or post-flight samples. These taxa were further filtered for taxa that were observed in at least two ISS samples. The resulting list included five taxa: two viral genera, two viral species (both phage), and one bacterial species: *Rickettsia australis* (Figure 2). Given the generally low abundance of these taxa, we cannot definitively rule out that they were present at an undetectable, low threshold pre-flight, but our data show no reads supporting their presence.

**Figure 2:**
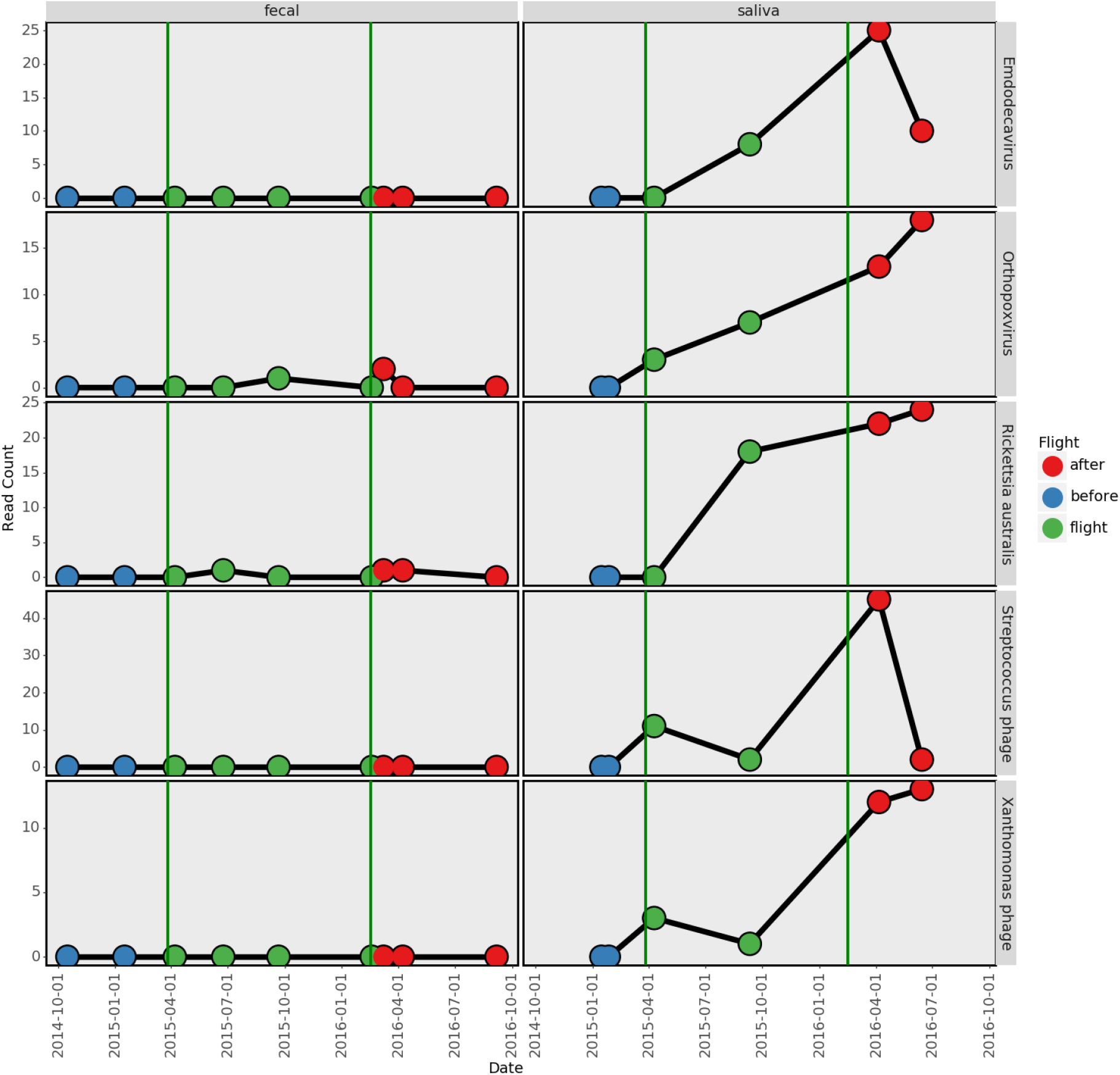
Total number of reads observed in TW for different taxa not observed before flight. Green vertical bars indicate the start and end of flight. The *Streptococcus phage* referenced is *phiARI0004, Xanthomonas phage* is *vB XveM DIBBI*,

#### Emergence of new taxa in gut microbiomes exceeds repeated sampling

To place these taxonomic trends in context, we investigated whether the sampling time series from TW and HR would identify more new taxa than repeated assays on an unchanging fecal sample. We compared the fecal microbiome time series of TW and HR to 243 repeated samples taken from a single fecal sample (Sasada et al., 2020), using 100,000 random sets of 9 samples (to match the twins’ set size). The number of taxa in each sample that had not been observed in any previous sample were counted for each subset and normalized by the total number of taxa in the first sample. The time series for HR showed significantly more new taxa than 99,971 (p = 2.9e-4) random stool subsets, and TW was even more significant (more than 99,990, p = 1.0e-4) relative to random subsets (Figure 3). These results shows that the time series for TW and HR both consistently had more taxa than would be expected from re-sampling an unchanged fecal sample. Differences between TW and HR may have a large number of causes including: diet, environment, and exposure to other people. For our subsequent analysis, it is relevant that both TW and HR show more new taxa than re-sampling a single sample, since it implies transmission may be occurring (possible for both subjects though we do not have data on HR’s environment for direct comparison).

**Figure 3:**
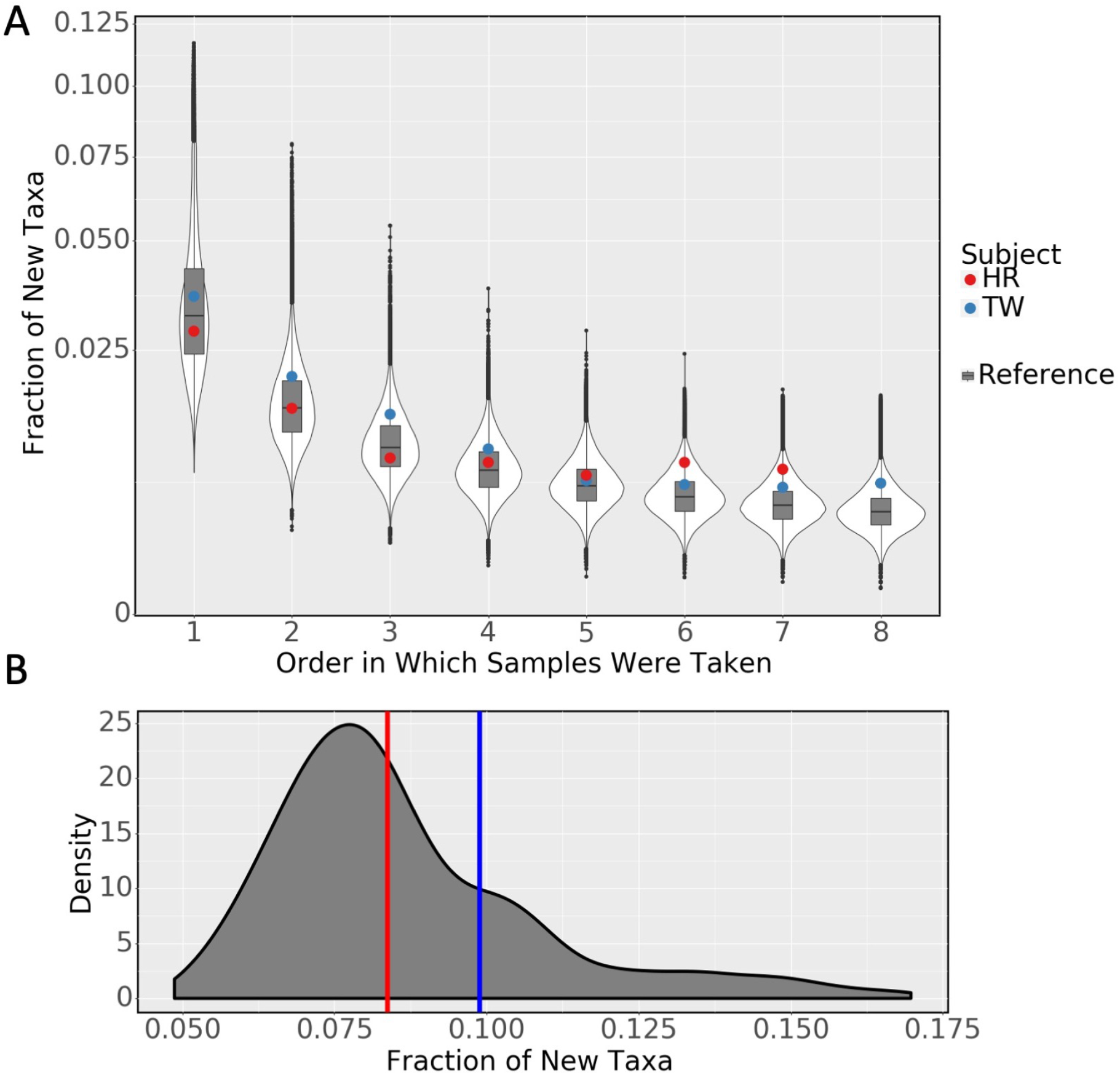
A) The number of new taxa observed in TW and HR are higher than repeated resampling of the same fecal sample. The y-axis gives the number of new taxa at each time point (not observed at any previous time point) divided by the number of taxa in the first sample. The first time point is omitted from the plot because it is always 1 by construction. The x-axis gives the order of each sample (arbitrary for random subsample). Boxplots show the distribution of random subsamples. Colored points are the actual time series. B) The number of unique taxa observed after the first time point divided by the number of taxa at the first time point. Same legend as (A)

#### Evidence of higher transfer rates on board the ISS

To characterize this possible microbial exchange, we next calculated taxonomic diversity using Shannon’s entropy for species profiles of each sample (Figure S2). For both fecal and saliva samples from TW, the highest diversity was observed during flight, and this trend was not observed for HR in the same time intervals. However, given the small sample size this trend was not significant (p=0.21). Nonetheless, we identified a significant increase in the number of previously unobserved taxa for samples taken from TW during flight (Figure S3) compared to random permutations.

To further characterize the significance of such transfer relative to the sampling set and the source, we performed a series of permutation tests. We first established the number of previously unobserved species found at each time point in the actual data from TW. We then randomly shuffled and relabeled these samples and counted species again for a total of 10,000 random permutations. We then counted the number of permutations where the number of species observed ‘during flight’ in the shuffled data was higher than the real data. For the fecal microbiome the actual number of observed taxa was higher than the shuffled data in 96.7% of cases, for saliva 98.2% of cases and for buccal the observed was higher than all other permutations. Repeating the same procedure on data from HR (‘flight’ status was arbitrarily assigned to the second, third, and fourth samples) we observed 45.9% for feces 98.2% for saliva, and 80.1% for buccal (more buccal samples were available for HR).

Results were similar when the above procedure was repeated only with taxa found in ISS environmental samples (TW fecal 97.5%, TW saliva 97.7%, TW buccal all permutations, HR fecal 33.6%, HR saliva 98.3%, HR buccal 81.1%). An analogous analysis performed on ISS samples (Figure S4) showed that microbial data during TW’s flight did not have significantly more new taxa than shuffled time periods (higher than 557 of 1,000 permutations). This is expected since the ISS is under continual habitation and merely is meant to show the converse of HR as a second control.

### 2.2 Strain level variation confirms microbial transfer

#### Novel genome regions in flight found in environmental and commensal microbiomes

Given the higher overall transfer rate of species on the ISS, we next examined the strain emergence and persistence (post-flight) of such species. We selected a set of candidate taxa that showed significantly greater abundance during and after flight in TW compared to pre-flight, and then mapped reads to known reference genomes from these taxa. We looked at the coverage of reference genomes in samples from TW at each stage of flight (concatenating samples from the same stage) and in the ISS and grouped regions into three categories: regions which were covered before flight, regions that were covered before flight in either gut or saliva samples but not observed in the other until flight, and regions that were not observed in either gut or saliva samples until flight but were found in the ISS environment. Example coverage plots are shown for two taxa: *Fusobacterium necrophorum* and *Serratia proteamaculans* (Figure S5 and Figure S6B respectively). The total size of these genomic regions for all tested taxa are listed in Table 2.

**Table 2:** Size of regions that may have been transferred in kilobases. Gut-Saliva transfer means that a region was found in either the gut or saliva microbiome pre-flight, then found in the other during-flight. Environment transfer means a region was not found in either fecal or saliva microbiomes from TW pre-flight but was found during flight and was also present in the ISS.

|  | Pre-flight | Gut-Saliva transfer | Environment transfer |
| --- | --- | --- | --- |
| *Bifidobacterium pseudocatenulatum* | 243.9 | 92.4 | 85.2 |
| *Brevibacterium siliguriense* | 18.7 | 2.6 | 3.1 |
| *Gordonibacter urolithinfaciens* | 37.8 | 12.9 | 21.2 |
| *Bacillus albus* | 87.6 | 7.0 | 14.1 |
| *Gluconobacter albidus* | 10.2 | 2.5 | 1.3 |
| *Fusobacterium necrophorum* | 86.4 | 18.0 | 56.8 |
| *Geobacillus stearothermophilus* | 73.5 | 13.7 | 13.8 |
| *Bifidobacterium catenulatum* | 258.8 | 17.5 | 40.3 |
| *Streptococcus viridans* | 2319.6 | 92.9 | 221.8 |
| *Vibrio alginolyticus* | 211.0 | 10.6 | 89.7 |
| *Staphylococcus sciuri* | 179.0 | 19.4 | 37.4 |
| *Pectobacterium parmentieri* | 269.0 | 22.7 | 56.7 |
| *Campylobacter lari* | 42.0 | 8.7 | 18.1 |
| *Atlantibacter hermannii* | 66.4 | 15.7 | 30.6 |
| *Bacillus tequilensis* | 57.4 | 6.0 | 8.7 |
| *Achromobacter ruhlandii* | 49.8 | 13.6 | 11.7 |
| *Serratia proteamaculans* | 70.0 | 11.2 | 6.7 |
| *Leptotrichia hongkongensis* | 115.2 | 0.7 | 21.5 |
| *Exiguobacterium antarcticum* | 21.5 | 4.4 | 6.2 |
| *Anoxybacillus amylolyticus* | 11.5 | 2.2 | 2.3 |
| *Kosakonia sacchari* | 65.4 | 16.0 | 30.8 |
| *Yersinia canariae* | 18.2 | 8.2 | 7.7 |
| *Providencia heimbachae* | 76.0 | 12.1 | 6.5 |
| *Spirochaeta perfilievii* | 2.7 | 0.4 | 0.7 |
| *Cronobacter condimenti* | 15.2 | 8.5 | 18.8 |
| *Brenneria rubrifaciens* | 13.2 | 5.7 | 7.3 |
| *Staphylococcus simiae* | 20.8 | 1.5 | 6.2 |

For the selected taxa, the average environmental transfer of genomic regions were 32.2% of the size of pre-flight regions, whereas gut-saliva transfers were lower at 19.9%. The taxa with the (proportionally) largest transferred regions *Cronobacter condimenti*, had 55.9% gut-saliva transfer and 123.7% environmental transfer. The presence of (in some taxa) large genomic regions that were not covered until flight strongly suggests that individual species are undergoing flux with new strains and genes migrating into commensal microbiomes and/or a greater abundance of the strain.

#### Microbial SNPs match environmental and commensal microbiomes

Once the candidate genomic regions were identified, we next mapped co-occurring clusters of SNPs (haplotypes) in the selected taxa listed above in all samples from TW, HR, and the ISS (Figure 4). We matched microbial haplotypes from TW during flight to possible sources in pre-flight TW samples and ISS samples. We considered five groups as potential sources for haplotypes in mid-flight fecal samples: haplotypes found in pre-flight fecal samples, haplotypes found in pre-flight saliva but not fecal or ISS samples, haplotypes found in the ISS but neither saliva nor fecal, haplotypes found in both the ISS and saliva (mixed) but not fecal, and haplotypes not observed in any other group. For saliva samples we used the same five groups but replaced fecal with saliva and vice versa.

**Figure 4:**
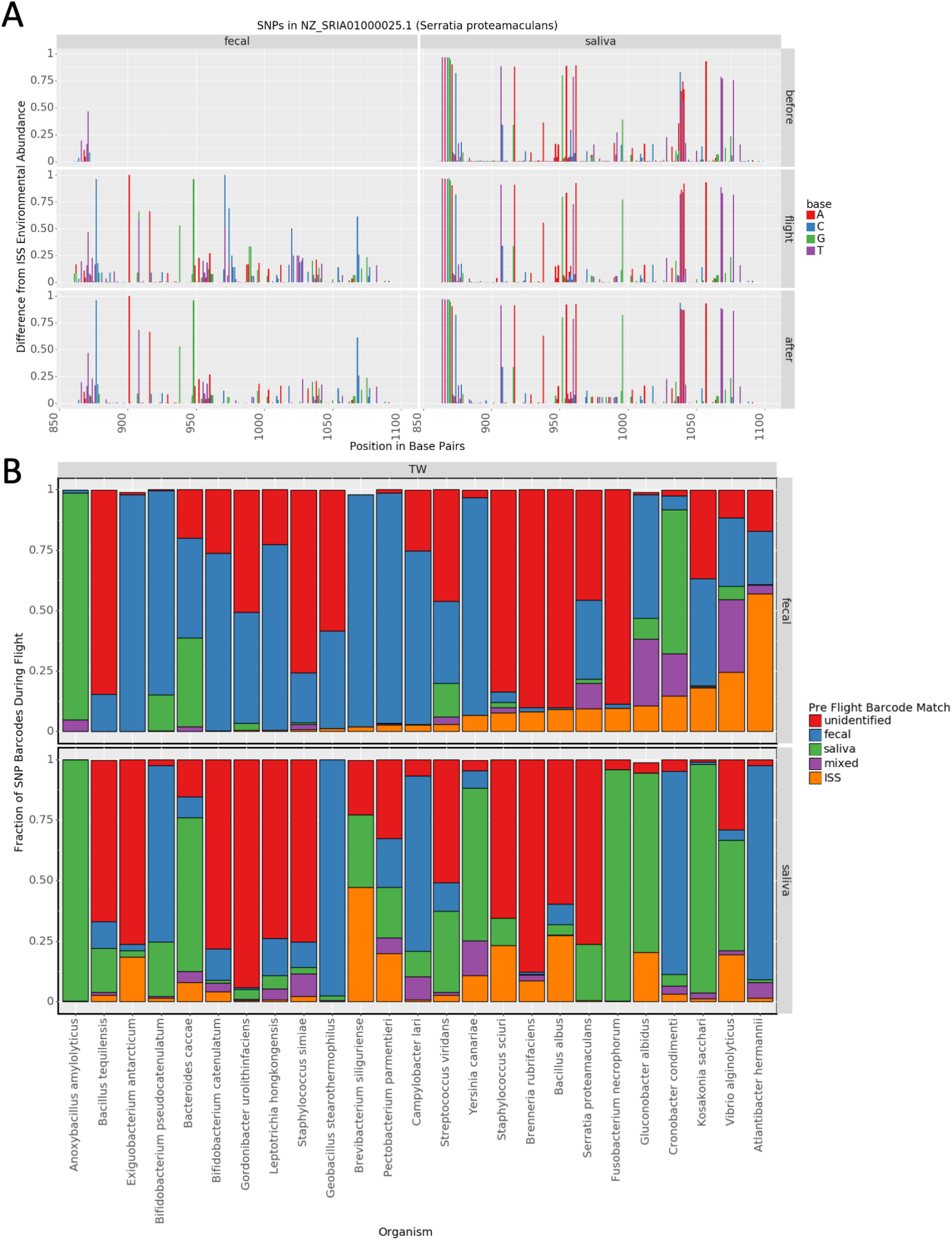
A) An example set of SNPs found in Serratia proteamaculans. The abundance of each SNP is shown relative to the frequency of the base found in the ISS at each position. A tall column indicates a base was low abundance in the ISS environment. In this case the SNPs shown for the fecal (left) strain match a secondary strain in the environment and constitute a candidate for transfer from the environment to the gut microbiome. B) Pre-flight sources of different SNP barcodes observed in TW during flight. Each SNP barcode in peri-flight samples from TW was matched to barcodes in pre-flight samples from TW and ISS samples. The fraction of barcodes matching each source is shown. For fecal samples barcodes labeled as saliva did not match fecal samples and vice versa. Barcodes labeled as matching ISS were not found in either fecal or saliva samples.

The pre-flight sources of haplotypes varied by the species being investigated (Figure 4B). Some species, such as *Cronobacter condimenti* showed an apparent flip of strains from the gut microbiome to saliva and vice versa. Other taxa, like *Atlantibacter hermannii*, showed a large fraction of haplotypes that matched environmental haplotypes in the gut microbiome. Some taxa, like *Bifidobacterium catenulatum* showed little similarity to any potential external source.

### 2.3 Transfer of *Serratia proteamaculans*

#### *Serratia proteamaculans* (SP) is a candidate of persistent transfer

We identified S. proteamaculans as a candidate species for persistent transfer, since it was found in ISS environmental samples and was significantly more abundant in mid- and post flight fecal samples from TW than in fecal samples from TW pre-flight and HR samples. Overall, S. proteamaculans was only found at low levels in fecal samples in TW pre-flight, was significantly more abundant during flight, and dropped to an intermediate level after flight (Figure S6A). No major variation in abundance was observed for the control twin HR. SP was roughly uniformly abundant in the saliva before during and after flight.

#### Regions of the SP genome are found in TW fecal samples only after arrival at the ISS

We identified regions of the SP genome which appeared in fecal samples after TW was on board the ISS. We found three such regions totaling about 1.5kbp. The abundance of these regions roughly matched the overall pattern seen for S. proteamaculans: very low or undetectable pre-flight, a high during flight, and an intermediate level post flight (Figure S6B). These regions were all well covered from ISS environmental samples.

Total coverage of the SP genome in TW from all available fecal samples was 29.2kbp. Before flight 8.9kbp was covered, during 17.2kbp and after 19.0kbp. However some of these regions were either quite small or not covered in both mid- and post flight. As, such 1.5kbp represents a reasonable fraction of the amount of SP genome covered in TW but should only be interpreted as evidence for the transfer of particular genes.

#### SNPs in post-arrival regions match a secondary environmental strain

We next analyzed one of the above regions (of about 250bp) for SNPs (Figure 4A) and identified SNPs in samples from TW which did not match the dominant strain in the ISS environment. We identified 9 such SNPs in this region in fecal samples taken from TW during flight. Of these 9 SNPs, 6 were also found after the conclusion of flight. No SNPs were found in fecal samples from TW before flight owing to the fact that no reads mapped to this region. We note that all 9 SNPs were found in the ISS environment at some proportion and that this region did not match any other reference genome in RefSeq besides SP.

Next we sought to determine if these 9 SNPs could have come from a secondary strain on the ISS. We used the SNP clustering technique described in the methods to determine if the 9 SNPs we identified could have come from the same underlying sequence. We identified corresponding barcodes (groups of co-stranded SNPs) of 8 SNPs in TW and 9 SNPs in the ISS environment. The groups in TW included 8 out of the 9 mid-flight SNPs. The 9 SNP group from the ISS environment included these same 8 SNPs as well as one SNP not identified in TW. This leads us to the conclusion that the strain found in TW likely represented a secondary strain in the ISS environment.

Finally, we checked if the strain could have come from TW’s oral microbiome. The region in question was covered by reads in the oral microbiome before flight. However, when we performed the same analysis as above using reads from TW’s oral microbiome we found a distinct SNP pattern (Figure 4A) giving evidence that the strain found in flight likely did not come from the oral microbiome.

### 2.4 Changes to immune repertoire in flight suggest environmental transfer

#### Immune repertoire in TW changes during flight

We surveyed the alpha (intra-sample) and beta (inter-sample) diversity for the repertoires of T-Cell Exposed Motifs (TCEMs) found in each sample from RNA-sequencing of CD4+ sorted peripheral blood mononuclear cells (PBMCs) from TW and HR (Garrett-Bakelman, 2019). To asses beta diversity we performed a UMAP dimensionality reduction using Manhattan distance between the TCEM repertoires (with abundance) from each sample. As expected, there was little similarity between the three types of TCEM motifs (types 1, 2a, and 2b Figure S7). Within each type of motif, there was weak clustering between the different stages of flight for TW and overlap with HR (Figure 5A). This indicates a possible shift during flight. The clustering does not seem to be due to a batch effect from the return of 3 flight samples directly from the ISS, as these samples cluster more closely with fresh frozen samples than they do with the ambient return sample from HR (Figure S8).

**Figure 5:**
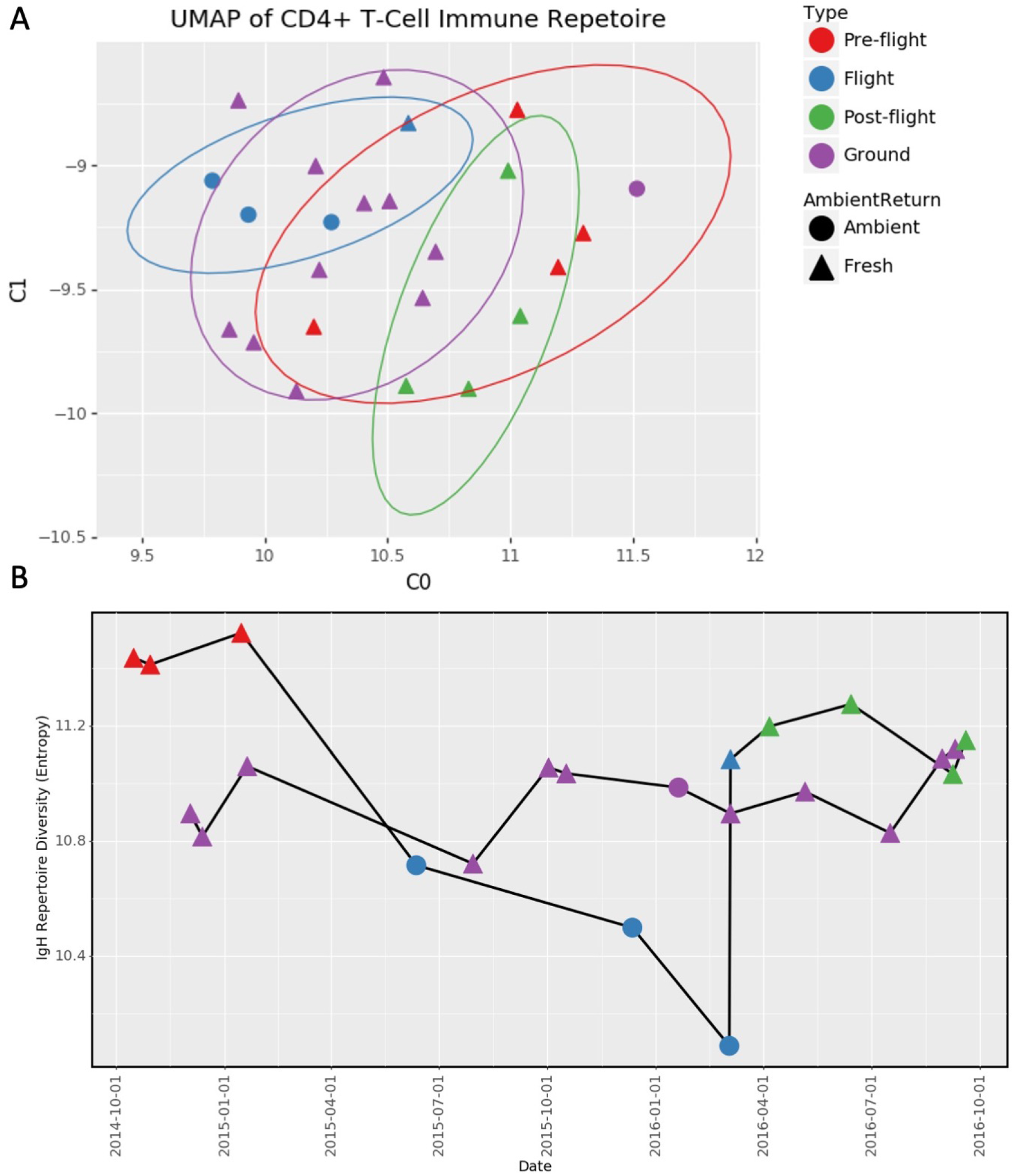
A) UMAP plot of similarity between Type 1 TCEM repertoires. B) Entropy of depth normalized TCEM repertoires. Legened is the same for both panels, ‘Ground’ indicates HR.

We next assessed alpha diversity by randomly subsetting TCEM repertoires from each sample to 5,000 total TCEMs (fewer unique TCEMs as the same sequence could be taken multiple times) and taking Shannon’s Entropy (*H* = −∑_*i*_ *p_i_*log_2_(*p_i_*)) of each sample (Figure 5B). Shannon’s entropy was used because it accounts for differences in abundance. We observed a large drop in diversity in TW during flight which partially recovered after flight. By comparison, the TCEM profile for HR was relatively stable (10.8-11.1).

#### TCEMs match environmental targets in the ISS

We next compared the overlap of TCEM repertoires in each sample to potential TCEM targets in ISS environmental samples. All PMA treated samples (to select for intact cell membranes, and likely bacterial viability) from the ISS during TWs flights were pooled. We produced a set of potential TCEM targets and filtered all targets that occurred in less than one part per million. This left a set of 176,204 potential environmental TCEM targets. We found the fraction of TCEMs (of all types) that occurred in the set of potential targets for each time sample from TW and HR. The fraction of TCEMs that overlapped with ISS targets increased in TW during flight (Figure 6A) and returned to an intermediate level after flight, and was the highest at the later points of the year-long mission. No corresponding change was observed in samples from HR.

**Figure 6:**
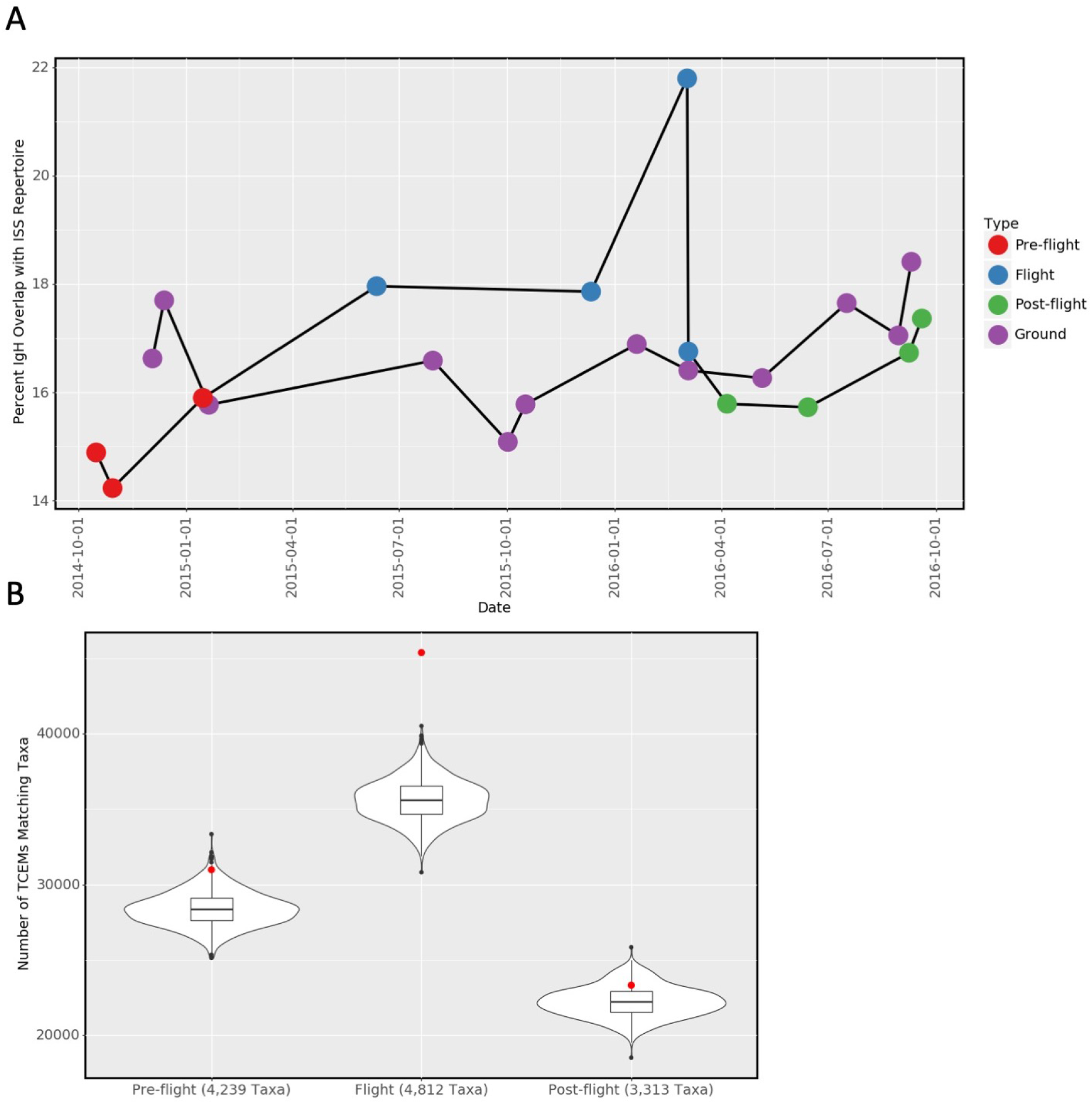
A) Overlap between TCEM repertoires and TCEM-targets found onboard the ISS. ‘Ground’ indicates HR. B) Number of TCEMs that match taxa found in both the ISS and flight (or pre and post-flight samples). The red dots indicate the actual values for each time point, the black distribution is made up of 10,000 random subsets of the same number of taxa for each time point. Many more TCEMs match ISS taxa from in-flight samples than pre-flight, post-flight, or random samples.

For further comparison, we took the anti-sense protein (as described in Root-Bernstein (2016)) of each TCEM and compared the set of anti-TCEMs for each sample to the environmental targets. The anti-sense protein has no clear biological significance for TCEMs so this was meant as a negative control. As expected there was no clear pattern in the overlap between anti-TCEMS and environmental targets (Figure S9).

#### Taxa found in the ISS are enriched for sequences that match TCEMs in TW during flight

We next identified taxa that contained proteins which matched the sequences for TCEMs. Since TCEMs are just 5 amino acids long and can match many different taxa, we pooled TCEM repertoires from flight and pre-flight samples from TW and found matching taxa for all TCEMs in the pooled repertoires.

Note that the pooled repertoires were randomly subsampled to contain the same number of TCEMS. We assessed how many taxa matching the TCEM repertoires were also found in the ISS: 4,812 matching taxa were found from the in-flight repertoire compared to 4,239 for pre-flight and 3,313 for post-flight.

Finally, we counted the total number of matches between TCEMs and taxa which were found in the ISS each time period (red dots in Figure 6B: with 45,373 found in flight, 30,982 for pre-flight, and 23,316 for post-flight). For comparison, we counted the number of matches found in 10,000 random subsets of taxa for each time period, where the size of the subsets was the same as the number of ISS taxa matched for each period. We found that the pre-flight taxa set had more matches then 9,856 random subsets (empirical p-value = 0.0144) while post-flight had more matches than 8,527 random subsets (p = 0.1473). However, the in-flight taxa set had far more matches than any tested random subset (p=<0.0001). This is evidence that the match between TCEM repertoires in-flight and environmental targets in the ISS far exceeds what would be expected by chance alone.

## 3 Conclusion

We have identified genetic and immunological evidence of microbial transfer between the fecal and saliva microbiomes of an adult and between these microbiomes and their environment. These data, derived from shotgun metagenomics sequencing and TCEMs, demonstrate that non-pathogenic microbes from the environment can establish themselves in astronauts and suggests the possibility of ongoing microbial flux between humans and the unique ISS environment. Moreover, these provide candidate “ISS mobile” species and also enable a key estimate of the fraction of taxa that could be transferred from different sources of the body while in the spaceflight environment.

A number of open questions remain. We have made a first attempt to quantify the rate of transfer between different microbiomes and given an estimate for the total number of emergent species in a gut microbiome, which cannot be explained as the result of repeated sampling alone. However, these estimates necessarily suffer from the small sample sizes available in this study and the unusual situation under which the samples were taken. To conclusively establish the scope of microbial transfer will require broader studies targeting earth based environments, food varieties, and different, communities, as well as additional characterization using culture-based techniques. Nonetheless, the unusual nature of spaceflight provides a strongly controlled environment, making this a near-optimal model for studying microbial transfer.

The emergence of new taxa, while intriguing, must be placed into the context of expected stool sampling variation. To account for such sampling dynamics, we also conducted a rigorous re-sampling study. Our data showed that TW and HR had more newly observed taxa at some (but not all) of the time points relative to the 100,000 subset. Importantly, the number of new taxa that were observed in subsets dropped off quickly for later time points as the subsets reached saturation. Subsets generally showed an adversarial selection, wherein many new taxa at one time point would lead to fewer new taxa at later time points. The 243 fecal replicates had similar read counts to the time series from HR and TW, reducing a source of potential bias, but could also be examined in greater detail in future studies.

Of note, repeated sampling can identify low abundance species which were dropped out of previous samples and because different sample preparation techniques can yield different sets of taxa. A series of samples taken from a microbiome that is exchanging taxa with an external environment will have an additional source of new taxa. These taxa would not be identified in earlier samples because they were not present, and this is another source of variation that could be mapped and quantified for future missions (more sampling of more areas of the body and the ISS, and at greater depth).

The techniques for comparing immunological signatures in T-Cell repertoires to microbiomes are nascent. To our knowledge, this is the first study to compare T-Cell repertoires and microbiomes in humans using genetic data, but these techniques used may be limited in scope or accuracy. Though our results suggest a shift of the T-Cell repertoire in response to the new and unique environment of the ISS, we must temper this until these techniques can be proved and validated in other studies. Nonetheless, such metrics can lay the foundation for a strong potential link between a person’s T-cell dynamics and their environment.

Taken together, the matching genomic regions across 16 taxa, the host immunological data, and matching SNP haplotypes within the strains strongly supports the conclusion that novel taxa in preflight commensal microbiomes from TW could come from the environment or from other commensal microbiomes. The size of transferred regions and number of SNPs suggests that “taxa transfer” between commensal microbiomes occurs more frequently than they transfer from the environment to commensal microbiomes. However, these rates may prove to be anomalous for either TW, habitation in the ISS, or both, since non-pathogenic microbial exchange with the environment represents a significant unknown for its impact on human and astronaut health. Nevertheless, accurate quantification of microbial strains and their movements can lead to targeted interventions, shed light on the hygiene hypothesis (broadly and on the ISS), and help in planning for future missions and astronaut monitoring.

## 4 Availability and Access

All analysis and figure generating code may be found on GitHub at https://github.com/dcdanko/twins_iss_transfer. All results and raw data may be found on Pangea at https://pangea.gimmebio.com/sample-groups/62661efb-a433-4ae5-bcec-de704a80e217.

## 5 Author Contribution

DCD performed all bioinformatics analyses and defined the structure of the study with guidance from CEM. NS led the collection of samples from the ISS. DJB and CM prepared samples for sequencing. PJ, AK, MMC, GC, EA, coordinated sampling. FGB prepared samples for sequencing. SJG and MHV handled sample coordination, sequencing, collection, analysis. KV led coordination with NASA and led collection of samples on board the ISS. CEM led and conceived this study.

## 6 Acknowledgment

We would like to thank the Epigenomics Core Facility at Weill Cornell Medicine, the Scientific Computing Unit (SCU), XSEDE Supercomputing Resources, the Starr Cancer Consortium (I13-0052), the Vallee Foundation, the WorldQuant Foundation, The Pershing Square Sohn Cancer Research Alliance, NASA (NNX14AH50G, NNX17AB26G), the National Institutes of Health (R01AI151059), TRISH (NNX16AO69A:0107, NNX16AO69A:0061), the Bill and Melinda Gates Foundation (OPP1151054), and the Alfred P. Sloan Foundation (G-2015-13964).

Part of the research described in this article was carried out at the Jet Propulsion Laboratory, California Institute of Technology, under a contract with NASA. This research was funded by 2012 Space Biology project NNH12ZTT001N grant 19-12829-26 under task order NNN13D111T awarded to K.V., which also funded a postdoctoral fellowship for NKS.

## 7 Declaration of Interests

The authors declare they have no competing interests that impacted this study. CEM is co-founder of Biotia and Onegevity.

## 8 Methods

### 8.1 Experimental setup and samples

We analyzed 18 fecal samples from two human subjects (9 each) and 42 environmental samples from the ISS. All samples were assayed with 2×150bp DNA shotgun sequencing and analyzed as described below. Exact sample handling and processing is described in the supplementary methods.

Human fecal samples were taken from two identical twins TW and HR both astronauts who had previously been in space. During the study TW was sent on a roughly 1 year flight to the ISS while HR remained on earth and functioned as a control. For many parts of this study samples from TW are grouped into pre-flight, peri-flight, and post-flight groups. As much as practically possible samples from HR were handled in an identical manner to samples from TW.

We note that the sampling of the ISS was initially planned and designed separately from the sampling of the human subjects.

### 8.2 Sequencing

Samples from the human subject were extracted with a DNA extraction protocol adapted from the Maxwell RSC Buccal Swab DNA kit (Catalogue number AS1640: Promega Corporation, Madison WI). Briefly, 300 μl of lysis buffer and 30 μl of Proteinase K was mixed and added to each swab tube. Swab tubes were then incubated for 20 min at 56 C using a Thermo Fisher water bath, removed from the tubes, and fluid was transferred to well 1 of the Maxwell RSC Cartridge. The swab head was centrifuged using a ClickFit Microtube (Cat. # V4741), and extracted fluid was added to the corresponding well of Maxwell Cartridge, and eluted in 50 μl of provided elution buffer.

Extracted DNA was taken forward to the Nextera Flex protocol by Illumina. Briefly, 30 μl of extracted DNA was taken into library prep protocol and run with 12 cycles of PCR. Libraries were cleaned up with a left sided size selection, using a bead ratio of 0.8x. The right sided size selection was omitted. Libraries were then quantified using a Thermo Fisher Qubit Fluorometer and an Advanced Analytical Fragment Analyzer. Libraries were sequenced on an Illumina HiSeqPE 50 × 2 at the Weill Cornell Epigenomics Core.

Samples from the ISS were sequenced according to the protocol described in Singh et al. (2018).

### 8.3 Processing Short Read Sequencing Data

#### Preprocessing and Taxonomic Profiling

We processed raw reads from all samples into taxonomic profiles for each sample using the MetaSUB Core Analysis Pipeline (Danko and Mason, 2020). This includes a preprocessing stage that consists of AdapterRemoval (Schubert et al., 2016), Human sequence removal with Bowtie2 (Langmead and Steven L Salzberg, 2013), and read error correction using BayesHammer (Nikolenko et al., 2013). Subsequently reads were assigned to taxonomic groups using Kraken2 (Wood et al., 2019). We generated a table of read counts giving the number of reads assigned to each species for each sample.

#### Identification of candidate species for strain level analysis

We analyzed our table of species level read counts to identify candidate lists of *transient* and *persistent* transfer species. We held a transient species to be one that was transferred from the ISS into the astronaut only while the astronaut remained in the ISS and which was be cleared after return to earth. We held persistent species to be those that were transferred from the ISS to the astronaut which remained after return to earth.

We statistically analyzed our table of read counts using Aldex2 (Fernandes et al., 2013). Remaining samples (from astronauts) were split into two groups. The first group was the control group and consisted of all samples from TW before flight and all samples from HR at any point. The second group was the case group and consisted of all samples from TW during flight. Samples from TW after flight were assigned to the control group for analysis of transients and to the case group for analysis of persistents. Aldex2 was used to identify differentially abundant taxa between the two groups. We selected all taxa that were significantly (q < 0.05 by Welch’s t-test with Benjamini Hochberg correction) more abundant in the case group than in the control group. We then filtered these two list (persistent and transient) to include only species found in the ISS samples (minimum 10 reads in 25% of samples).

#### Strain Analysis

Reads were further processed for strain level analysis using the MetaSUB Core Analysis Pipeline. Given a specified organism to examine we downloaded all available reference genomes from RefSeq. If more than 100 reference genomes were available we selected 100 at random. Human-depleted reads were mapped to each genome using Bowtie2 (sensitive presets) and pileup files were generated using from alignments using samtools (Li et al., 2009). Pileups were analyzed for coverage patterns using purpose built code (see availability for access). SNPs were identified by comparing aligned bases from short reads to reference sequences, SNP filtering was performed as part of identifying co-stranded SNPs.

#### Identifying co-stranded SNPs

We developed a technique to identify SNPs that occurred on the same genetic strand. The technique is, in practice, limited to identifying co-stranded SNPs within 1kbp of on another. The technique works by formulating SNP recovery as an instance of the multi-community recovery problem. We start by building a graph of SNPs. Each SNP forms a node in the graph and is identified by its genomic position and base. Edges are added between SNPs that are found on the same read. Edges are undirected but weighted by the number of times a pair of SNPs is found on the same read. The SNP graph is then filtered to remove SNPs that occur only once as these are likely to be errors and are uninformative in any case. The remaining graph is clustered into groups of SNPs using the approach to the multi-community recovery problem by Blondel et al. (2008). The final result of this are sets of SNPs that are often found on the same read.

This technique is similar to techniques used for phasing SNPs to one strand of a diploid genome such as Zheng et al. (2016). The key difference between this technique and ours is that there may be more than two communities in our case and that we make only attempt to cluster proximal SNPs.

### 8.4 Analyzing Human and Environmental Immune Repertoires

#### Sample collection, preparation, and sequencing

Samples were collected and sequenced according to the protocal described in Garrett-Bakelman et al. (2019b). Briefly, PBMCs were flow sorted for CD4+ t-cells. RNA from these CD4+ T-Cells was selected using Poly-A pulldown leaving, largely, only RNA that would be translated and depleting ribosomal RNA. PolyA selected RNA was sequenced using Illumina machines for 2×150bp reads.

#### Assembling Immunoglobulin Heavy (IgH) sequences from short reads

We used MiXCR (v3.0.13) to build IgH sequences from poly-a selected RNA taken from CD4+ t-cells (Monk et al., 2017). We used the recommended workflow for non-specific RNA sequences and selected for IGH sequences using the built in export tool. The precise commands used are recorded in the attached *run_mixcr.py* file.

#### Creating a repertoire of t-cell exposed motifs (TCEMs) from IgH sequences

We assembled a repertoire of TCEMs from our IgH sequences following the method described by Breme and Homan (2015). Briefly, this method consists of taking 5 amino acid (aa) sub-sequences from IgH sequences. The 5aa sequences are found according to three specific spaced patterns within larger windows of 9, 15, and 15 amino acids respectively. These patterns are meant to reflect binding potential within the MHC groove. TCEM sequences are found by iterating every possible window (of both sizes) along the length of an IgH sequence and generating a 5aa sequence using the appropriate window. For a sample (or set of samples) we generated a full TCEM repertoire by concatenating the repertoire for each IgH sequence. For reference we note that there are only 20^5^ = 3.2 * 10^6^ possible 5aa sequences so significant overlap is possible between even randomly generated sets.

#### Creating a repertoire of TCEM targets from metagenomic data

We generated sets of potential amino acid sequences that could match our TCEM sequeces from metagenomic data. We counted all canonical 15 base pair nucleotide sequences from pre-processed and error corrected (see above) reads using Jellyfish (Marçais and Kingsford, 2011). We translated all resulting 15-mers and their reverse complement using the standard codon to aa table and discarded any aa sequence that contained a stop codon. Count information was retained for aa sequences from jellyfish. Any 5aa sequence that occurred with a frequency of less than 1 part per million was discarded.

#### Comparing metagenomic and t-cell repertoires

We compared the number of 5aa sequences which were found in both our metagenomic data and IgH sequences. Even with abundance filtering metagenomes typically still contained many more sequences than IgH repertoires. To asses overlap bewtween an IgH repertoire and a metagenome we took the number of 5aa sequences from the IgH repertoire which were also found in the metagenome divided by the total number of 5aa sequences in the IgH repertoire.

#### Identifying potential taxonomic matches to TCEM repertoires

We identified potential taxonomic matches to 5aa TCEM sequences by aligning them to the NCBI NR protein database using BLASTp. We accepted all 100% identity matches that included a taxonomic label. Since 5aa sequences are too short to be specific to one taxa there were typically multiple taxa for each TCEM. For a repertoire of TCEMs we built a binary matrix of TCEM sequences and taxa, each element in this matrix was set to true if and only if the given TCEm was found in the given taxon. This enabled us to generate statistics such as the number of TCEMs which could potentially match a taxon or the number of taxons for each TCEM.

## Supplement

**Figure S1:**
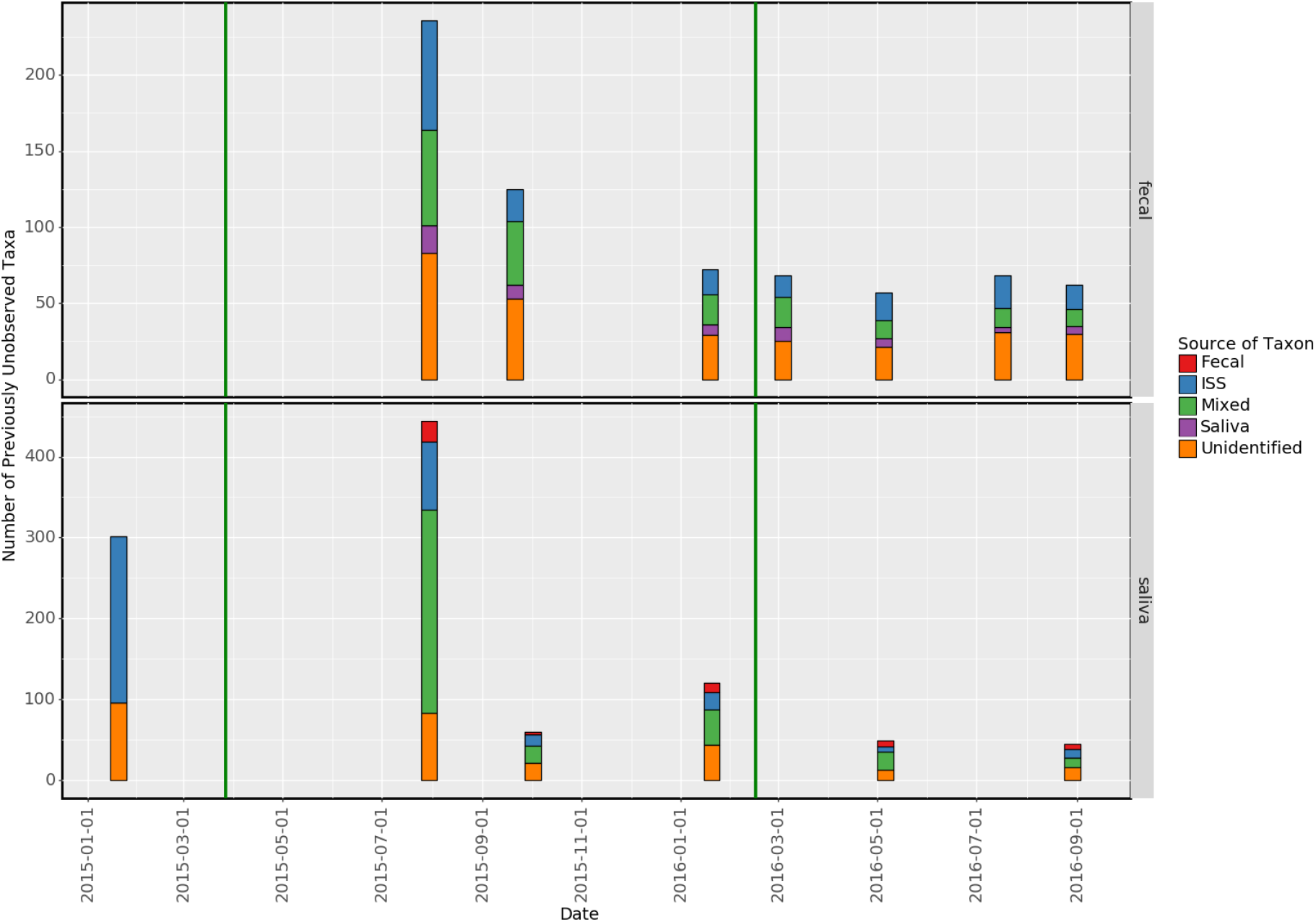
This plot shows the number of taxa at each time point that were not observed at any previous timepoint for fecal and saliva samples from HR. The colors indicate the likely source of the new taxon if it was found previously in the saliva (for fecal samples, vice versa for saliva samples), the ISS, both (Mixed), or neither.

**Figure S2:**
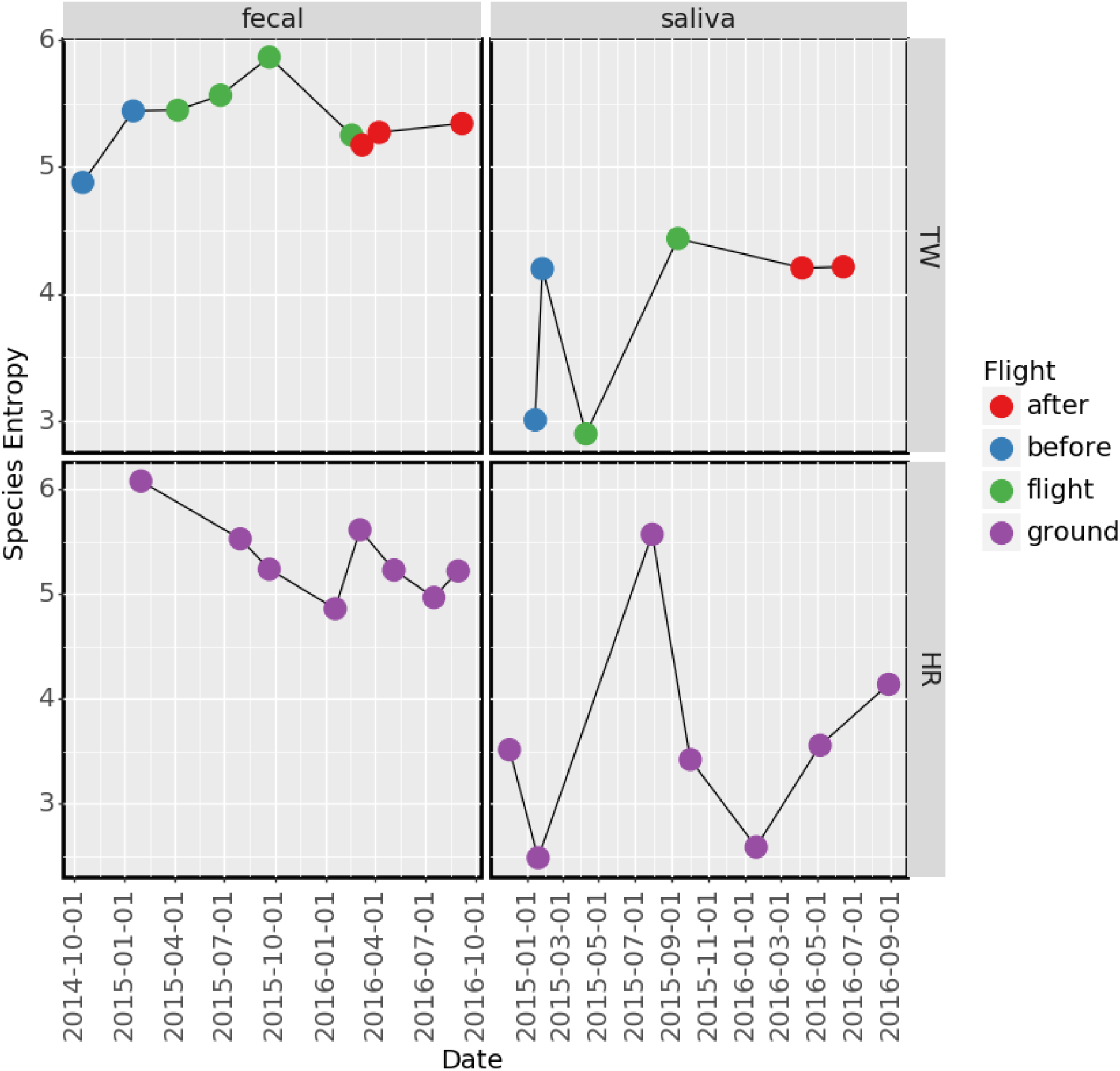
Vertical shows species entropy (Shannon entropy of species relative abundances) for sample types in both twins.

**Figure S3:**
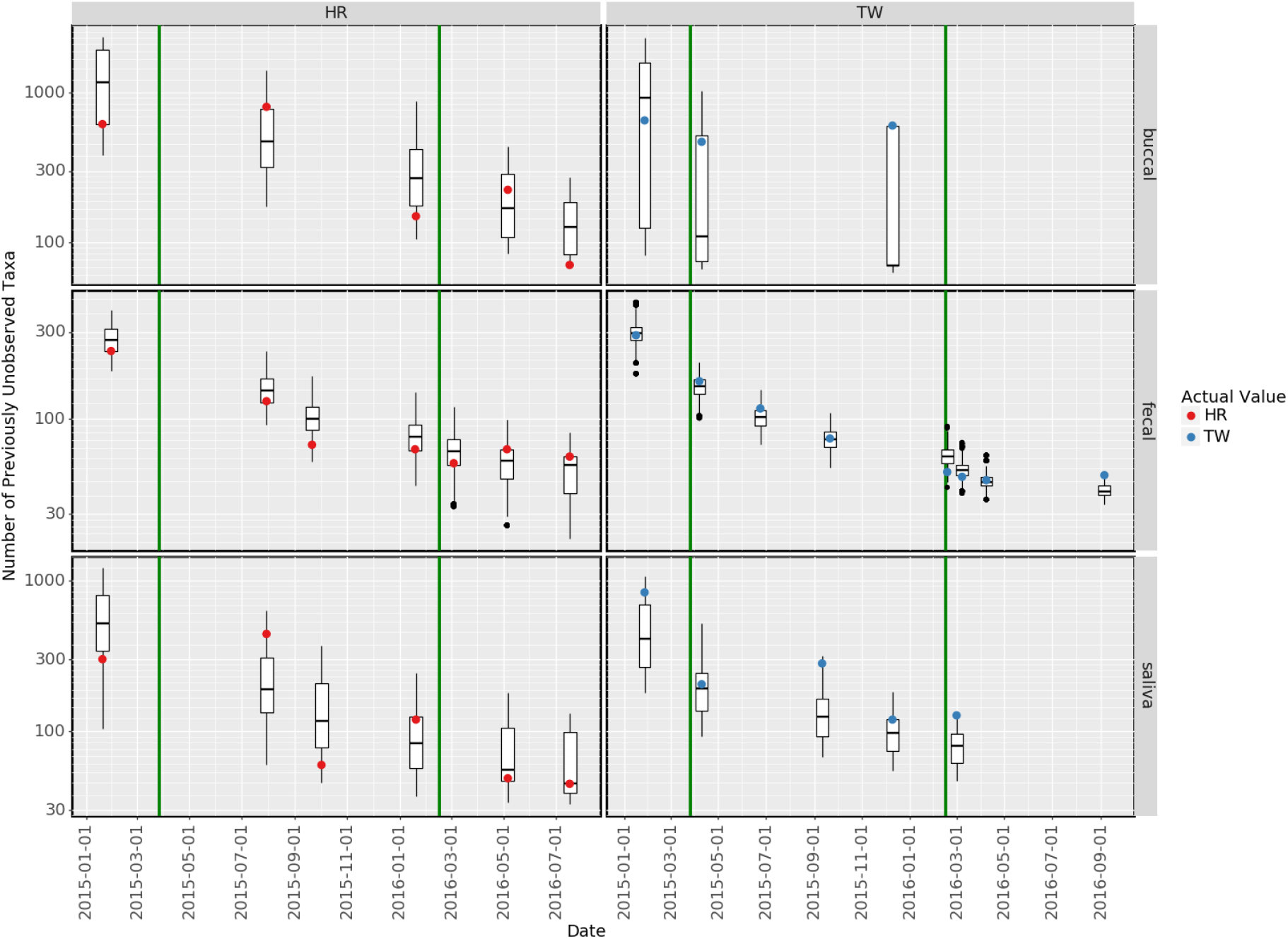
This plot shows the number of taxa at each time point that were not observed at any previous timepoint. The first timepoint is omitted from the plot since no taxa had been previously observed. Boxplots indicate an artificial reference distribution generated by randomly permuting timestamps. Red and blue dots indicate actual values.

**Figure S4:**
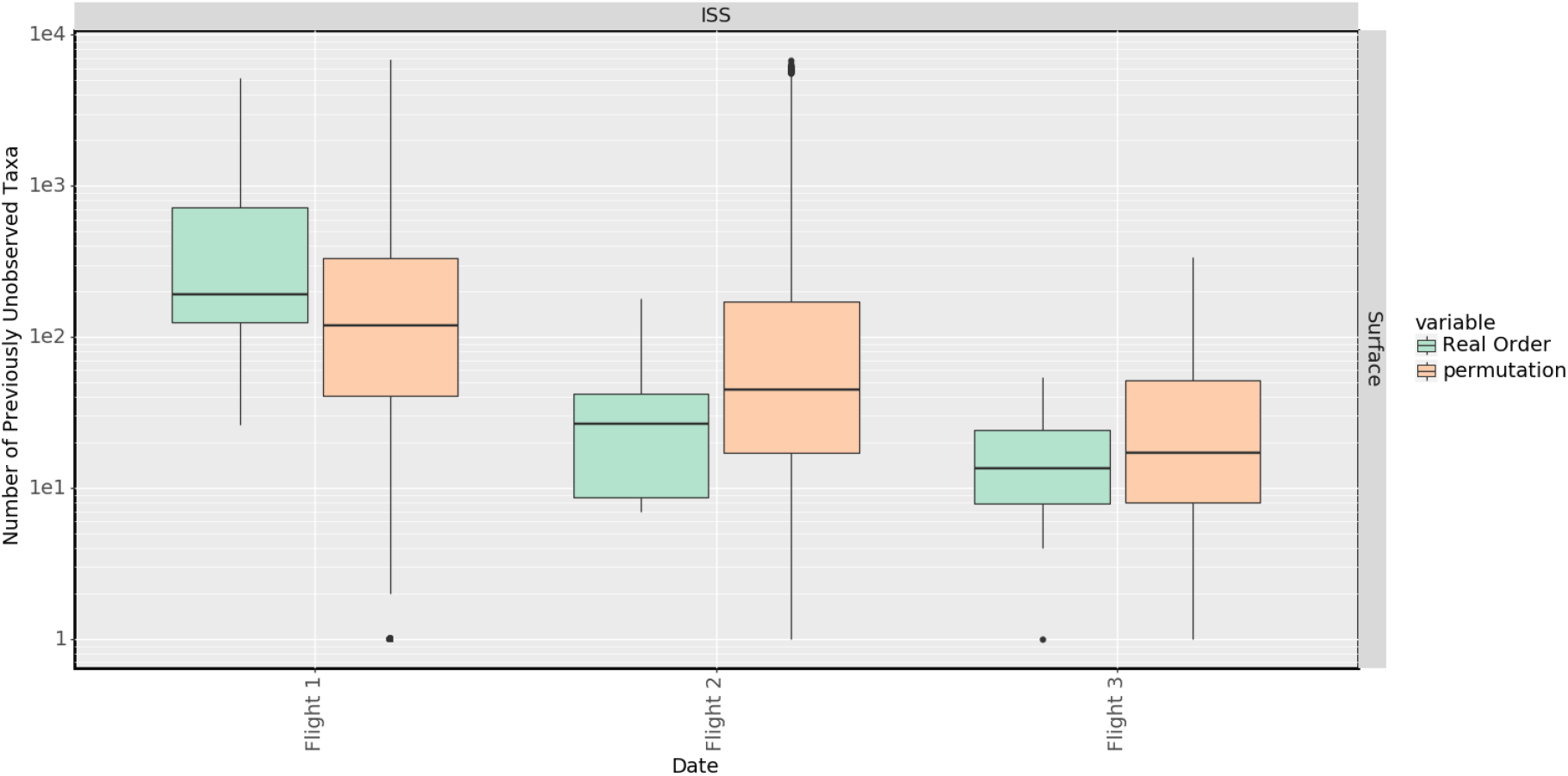
This plot shows the number of taxa at each time point that were not observed at any previous time point for the ISS. ISS samples are grouped into ‘flights’ where each sample in the same flight was taken on the same day. One sample from flight 1 is arbitrarily chose as the ‘first’ sample and used as the comparison. Boxplots indicate the real distribution of new taxa as well as an artificial reference distribution generated by randomly permuting timestamps.

**Figure S5:**
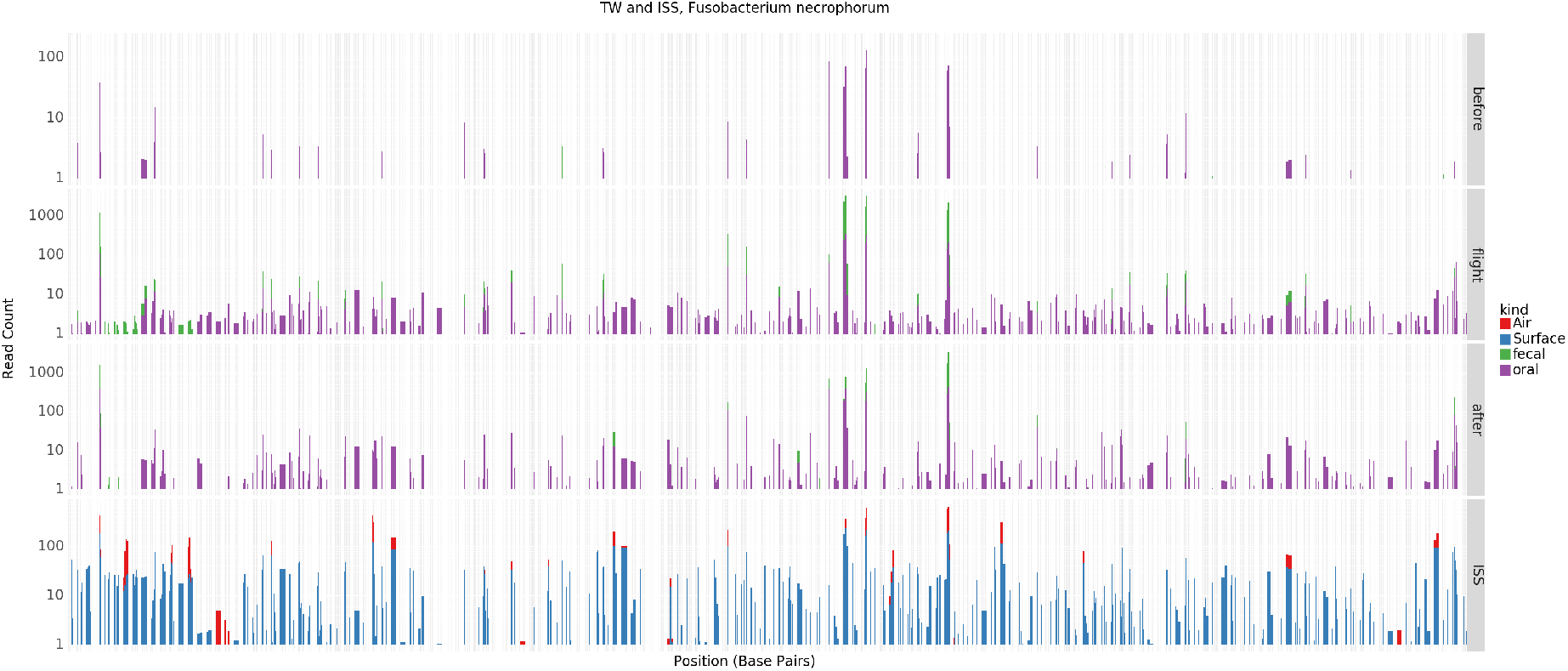
Rows show consolidated samples from before, during and after flight (or from the ISS at any point) from TW. Columns represent all available contigs for taxon. Colored bars represent 100bp covered, on average, at the specified read depth. A number of contigs are only covered in TW during and after flight.

**Figure S6:**
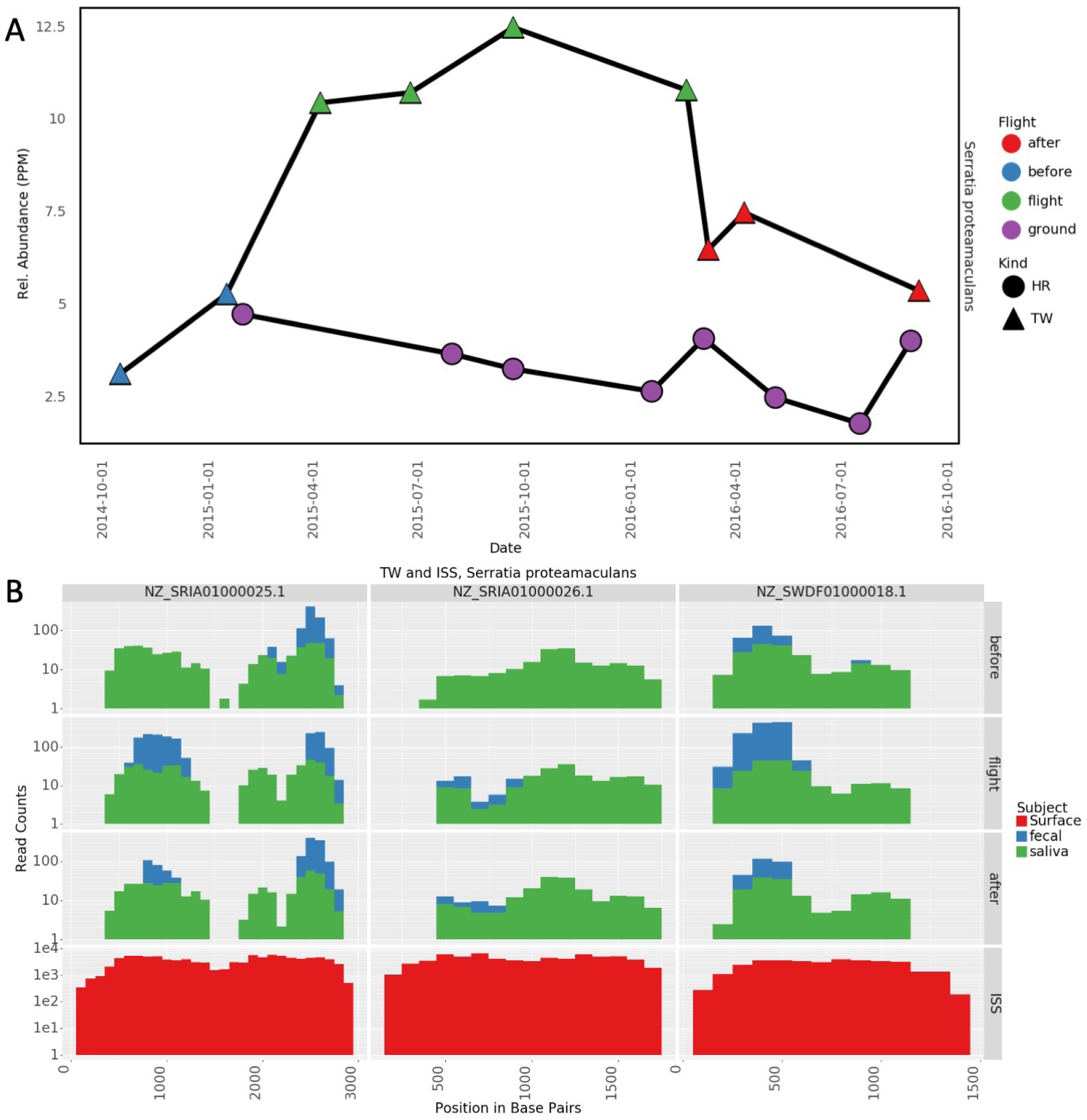
A) Relative abundance of *Serratia proteamaculans* in fecal samples from TW and HR. Relative abundance is given in units of parts per million. B) Coverage of candidate persistent transfer regions of the *Serratia proteamaculans* genome.

**Figure S7:**
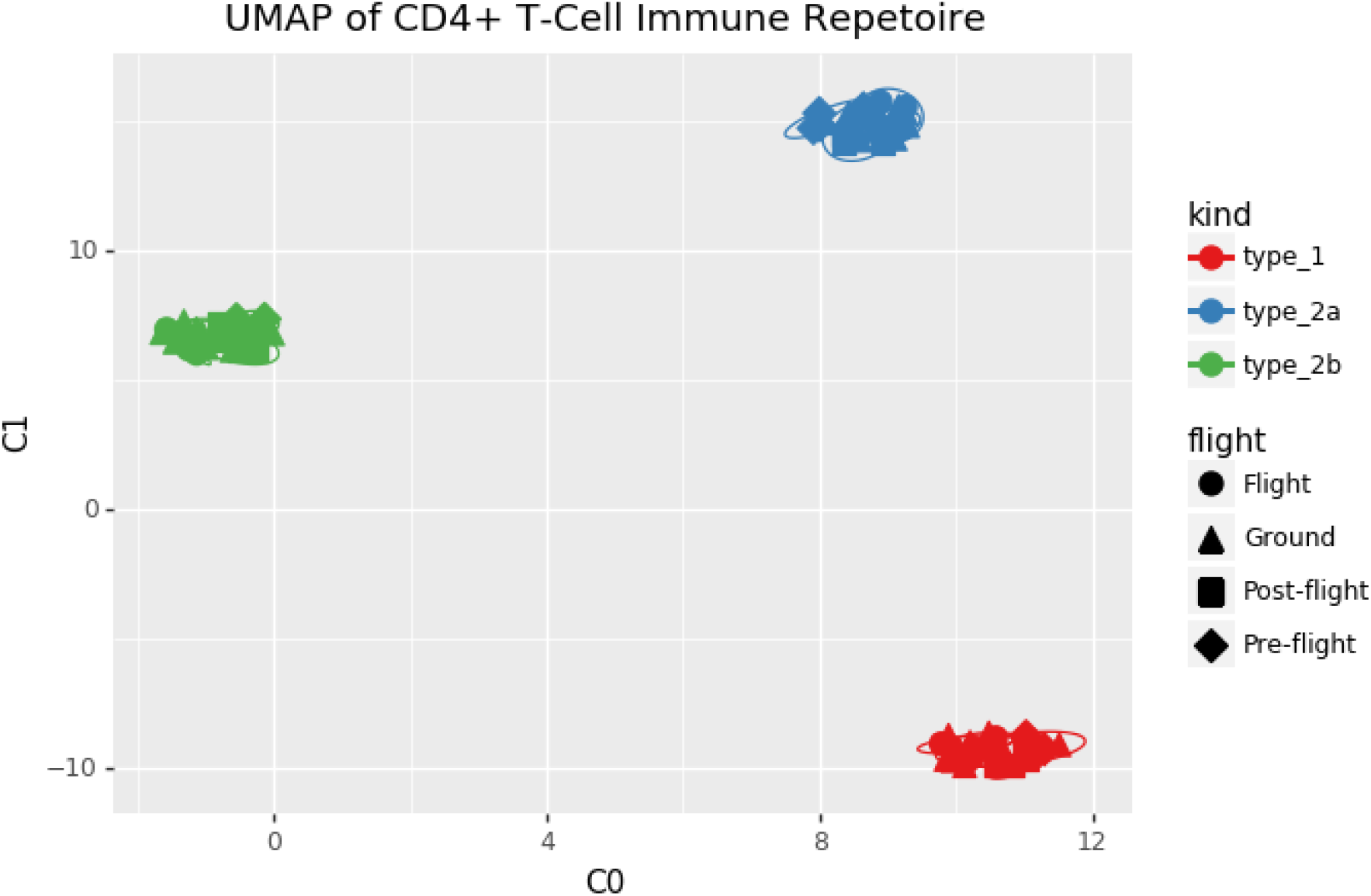
UMAP plot of similarity between TCEM repertoires of all types. ‘Ground’ indicates HR.

**Figure S8:**
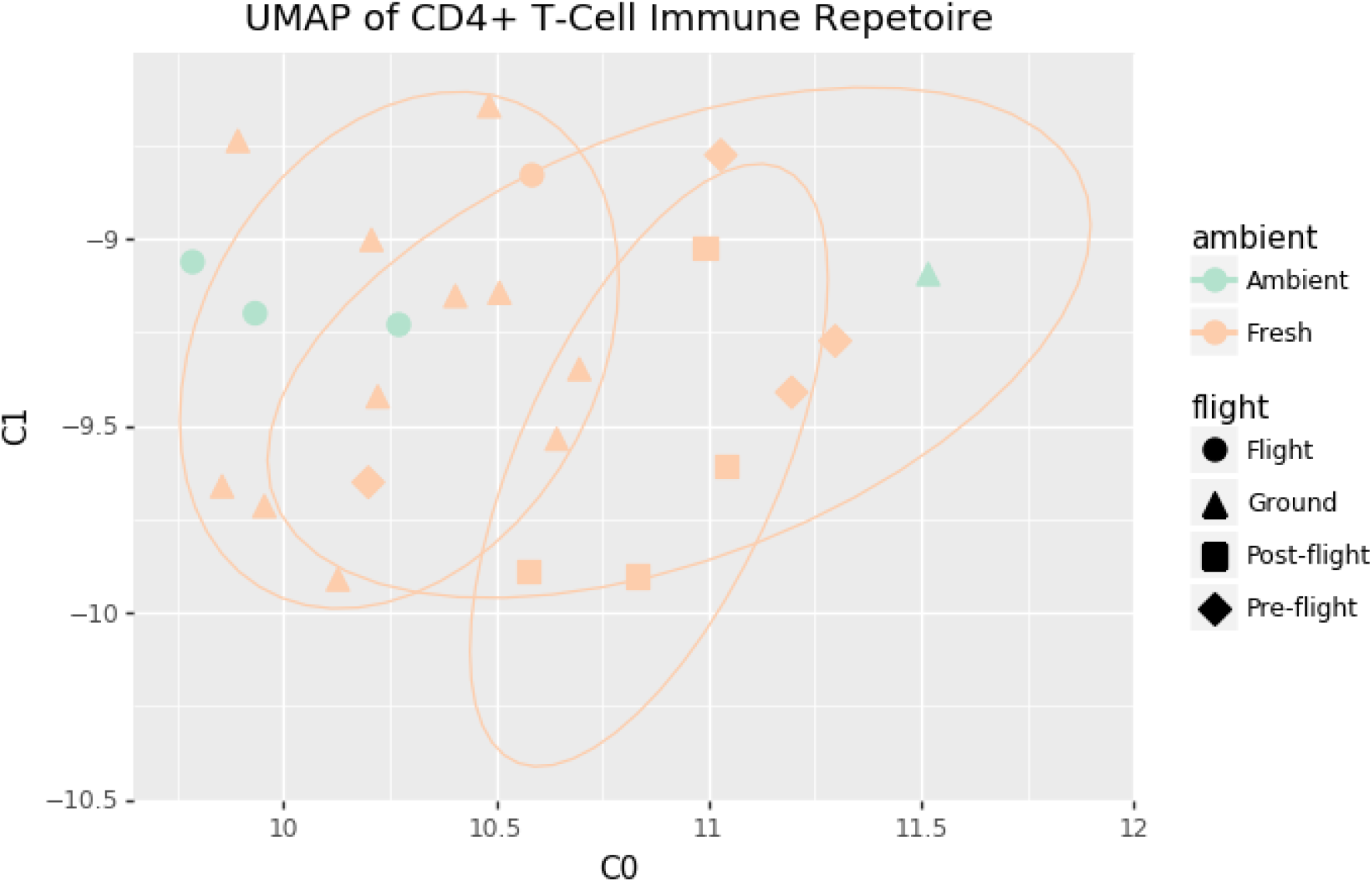
UMAP plot of similarity between type 1 TCEM repertoires of all types. Color shows method of retur

**Figure S9:**
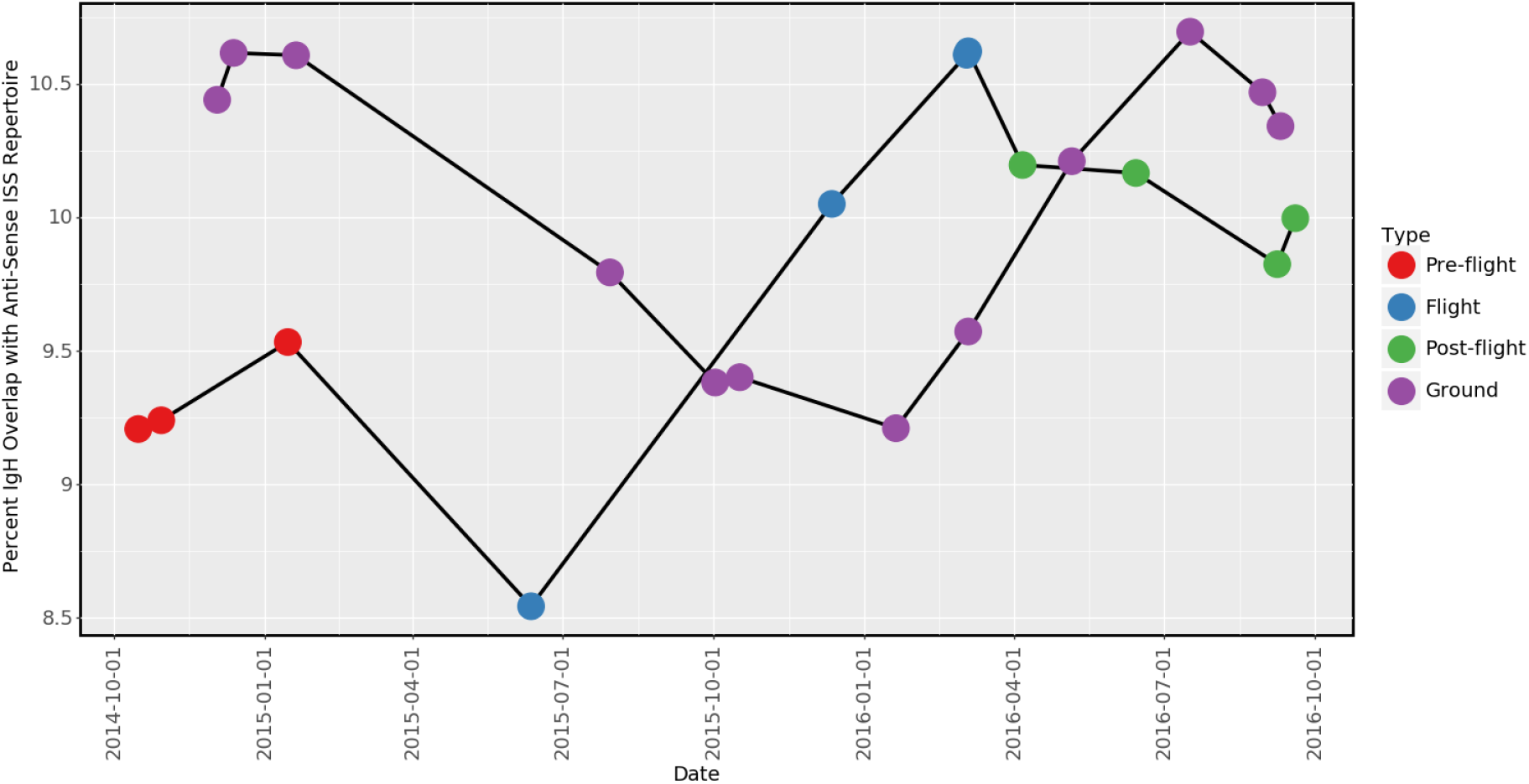
Overlap between anti-sense TCEM repertoires and regular TCEM-targets found on board the ISS. ‘Ground’ indicates HR. Anti-sense proteins are not expected to match, this serves as a negative control test.

## Notes

https://github.com/dcdanko/twins_iss_transfer

### Competing Interest Statement

The authors have declared no competing interest.

### Summary of Updates

this version of the paper includes a section relating the microbiome of the spacestation to an immune response in the astronaut

